# Protocol for Community-created Public MS/MS Reference Library Within the GNPS Infrastructure

**DOI:** 10.1101/804401

**Authors:** Fernando Vargas, Kelly C. Weldon, Nicole Sikora, Mingxun Wang, Zheng Zhang, Emily C. Gentry, Morgan W. Panitchpakdi, Mauricio Caraballo, Pieter C. Dorrestein, Alan K. Jarmusch

## Abstract

**Rationale:** A major hurdle in identifying chemicals in mass spectrometry experiments is the availability of MS/MS reference spectra in public databases. Currently, scientists purchase databases or use public databases such as GNPS. The MSMS-Chooser workflow empowers the creation of MS/MS reference spectra directly in the GNPS infrastructure.

**Methods:** An MSMS-Chooser sample template was completed with the required information and sequence tables were generated programmatically. Standards in methanol-water (1:1) solution (1 μM) were placed into wells individually. An LC-MS/MS system using data-dependent acquisition in positive and negative modes was used. Species that may be generated under typical ESI conditions are chosen. The MS/MS spectra and MSMS-Chooser sample template were subsequently uploaded to MSMS-Chooser in GNPS for automatic MS/MS spectral annotation.

**Results:** Data acquisition quickly and effectively collected MS/MS spectra. MSMS-Chooser was able to accurately annotate 99.2% of the manually validated MS/MS scans that were generated from the chemical standards. The output of MSMS-Chooser includes a table ready for inclusion in the GNPS library (after inspection) as well as the ability to directly launch searches via MASST. Altogether, the data acquisition, processing, and upload to GNPS took ~2 hours for our proof-of-concept results.

**Conclusions:** The MSMS-Chooser workflow enables the rapid data acquisition, analysis, and annotation of chemical standards, and uploads the MS/MS spectra to community-driven GNPS. MSMS-Chooser democratizes the creation of MS/MS reference spectra in GNPS which will improve annotation and strengthen the tools which use the annotation information.

## 1. Introduction

Identifying the chemicals detected in untargeted metabolomics experiments is challenging. Mass spectrometry (MS) is one method by which untargeted metabolomics can be performed. MS detects the mass of ions, *i.e.* mass-to-charge (*m/z*), as well as their abundance. Tandem MS (*aka* MS/MS), provides information about the molecular structure of chemicals. MS/MS, specifically product ion scans, contain interpretable patterns of product ions from which structural information is derived. Manual interpretation of MS/MS spectra is impracticable given the large amount of data generated in untargeted metabolomics experiments.

One solution to the identification challenge is to match MS/MS spectra with that of MS/MS reference spectra to provide an annotation. MS/MS matches to reference library spectra are considered level 2 or level 3 annotations based on the Metabolomics Standards Initiative (MSI).1 Reference libraries are time-consuming, costly to generate and often focused on a particular type of chemical class or the research interests of a laboratory or institute. Curatr, a web-based application for library generation, is one solution that improves the economy of effort.^2^ The Global Natural Products Social Molecular Networking (GNPS) platform^3^ offers a community-built MS/MS reference library which is part of the suite of libraries available for use, *e.g.* MassBank (Japan, European Union, and North America),^4,5^ ReSpect,^6^ MIADB, CASMI,^7^ PNNL lipids,^8^ Sirenas/Gates, and EMBL MCF. ^2^ While the option to contribute MS/MS spectra to the GNPS spectral library has existed since its inception, it remains a time-consuming process to select reference MS/MS and provide the required information for association with the entry (*e.g.* InChi, SMILES, CAS number, adduct, charge, and instrument).

To address the limitations in building public MS/MS spectral libraries, we have created a workflow directly into GNPS (dubbed MSMS-Chooser), borrowing concepts in Curatr to empower the GNPS community. The protocol details how to collect and contribute MS/MS spectra to the GNPS MS/MS reference library, including instructions for sample preparation information, a template for data entry, and web-enable data processing workflow in GNPS. To illustrate the utility of MSMS-Chooser, we generated MS/MS spectra from chemical standards in a 96-well plate format quickly, accurately, and with minimal manual interpretation.

## 2. Experimental

### 2.1 MSMS-Chooser Template

Chemical standards used to test the MSMS-Chooser workflow (**Figure 1A**) were purchased from Sigma-Aldrich (SKU: MSMLS-1EA). The MSMS-Chooser template (**Table S1**) was completed using the information provided with the purchased chemical standards. The MSMS-Chooser workflow uses the information in the MSMS-Chooser template to calculate the monoisotopic mass [M] and subsequently calculate the monoisotopic *m/z* of adducts. The current set of adducts considered are [M+H]^+^, [2M+H]^+^, [M+Na]^+^, [2M+Na]^+^, and [M−(H_2_O)_n_+H]^+^, [M−H]^−^, [2M−H]^−^, [2M−2H+Na]^−^; additional adducts can be added and retrospectively determined in data deposited in MassIVE, an MS data repository (massive.ucsd.edu). Additional information provided in the MSMS-Chooser template is used to automatically populate the fields required to upload MS/MS reference spectra to the GNPS library, such as the InChi and SMILES which are used to represent the chemical structure.

**Figure 1.**
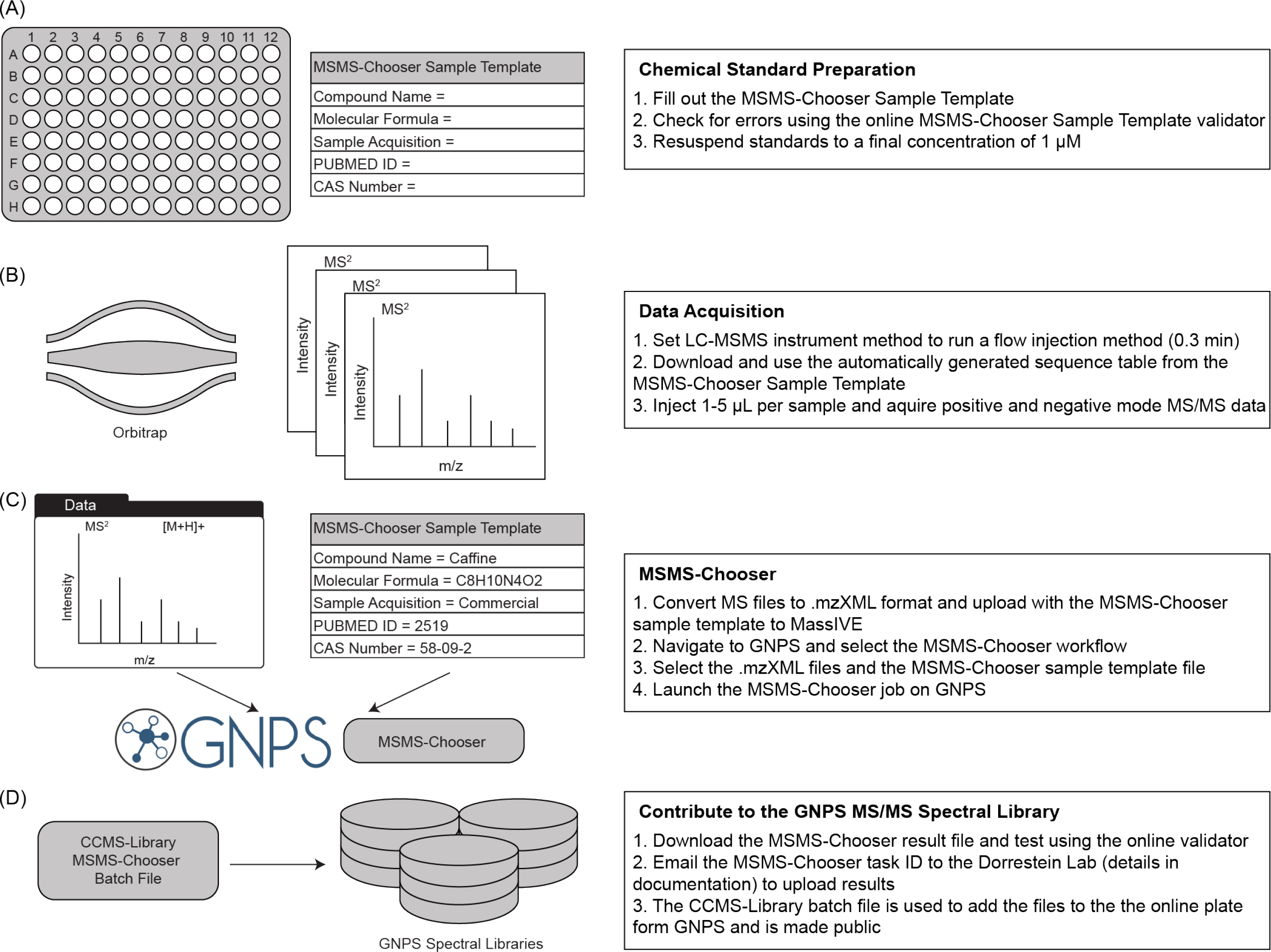
Overview of the MSMS-Chooser workflow and protocol.

### 2.2 Sample Preparation and Data Acquisition

Chemical standards were resuspended to a final concentration of 1 μM using a resuspension solvent of methanol-water (1:1) and transferred to a 96-well plate (**Figure 1A**). An ultra-high performance liquid chromatography (UHPLC) system (Vanquish, Thermo) coupled to an orbitrap mass spectrometer (QExactive, Thermo) was used (**Figure 1B**). The MSMS-Chooser template automatically populates a sequence table compatible with the Q-Exactive (future updates will include additional instrument specific sequence table templates). The data acquisition time was set to 0.3 min for each injection. Separate positive and negative ionization mode methods were used (*i.e.* one injection in positive mode and one injection in negative mode); the major differences in the methods are the ionization source parameters. 1 μL was injected from each well into a flow of solvent. The flow rate was set to 0.500 mL min^−1^ with a composition of 0.1% formic acid in water and 0.1 % formic acid in methanol (1:1). More detailed information on the instrument method and method files can be found in the MSMS-Chooser documentation (https://ccms-ucsd.github.io/GNPSDocumentation/msmschooser/).

### 2.3 Data Processing and Upload

Thermo’s proprietary MS file format (.raw) were converted to the open-source format (.mzXML), using the ProteoWizard tool MSConvert.^9^ All MS files (.raw and .mzXML files) and the MSMS-Chooser template were uploaded to MassIVE. It is recommended that all MSMS-Chooser datasets uploaded to MassIVE be prepended with the following text, “GNPS - MSMS-Chooser -”. Upon completion of the upload, the MassIVE accession was made public. Classyfire was used to generate chemical class information for the chemical standards by querying the SMILES.^10^

### 2.4 MSMS-Chooser

The MSMS-Chooser (v1.0) workflow, **Figure 1C**, is accessed on GNPS via the following link: https://gnps.ucsd.edu/ProteoSAFe/index.jsp?params=%7B%22workflow%22:%22MSMS-CHOOSER%22%7D with the source code available at https://github.com/CCMS-UCSD/GNPS_Workflows. The .mzXML/.mzML files (positive and negative mode) and the MSMS-Chooser template were selected into the “spectrum file” folder and “annotation table” folder respectively. Upon completion of the job, the resulting file output of MSMS-Chooser should be checked manually for accuracy, and poor quality or incorrect spectra should be deleted (row-wise) from the results table. The MSMS-Chooser output was then used to generate the associated MS/MS library spectra which were subsequently uploaded to the GNPS MS/MS spectral library (**Figure 1D**). Detailed instructions can be found in the MSMS-Chooser documentation (https://ccms-ucsd.github.io/GNPSDocumentation/msmschooser/). The MSMS-Chooser result page and resulting table are available via the following link https://gnps.ucsd.edu/ProteoSAFe/status.jsp?task=8454490b1ecc49ab85e1cece2f2f944c. A MASST search^11^ was performed on an illustrative compound, biotin, from the MSMS-Chooser results page (**Figure S1**). The illustrative MASST job can be accessed using the following link: https://gnps.ucsd.edu/ProteoSAFe/status.jsp?task=39cd886540c147e9a8a618b275e5f541.

### 2.5 Data Availability

QExactive tune files and instrument methods for positive and negative mode can be found on MassIVE via the following accession number: MSV000084286. The data (.raw and .mzXML) acquired for the Mass Spectrometry Metabolite Library (MSMLS) from Sigma-Aldrich and the associated MSMS-Chooser template can be found on MassIVE via the following accession number: MSV000084072. The MSMLS MSMS-Chooser job can be accessed via the following link: https://gnps.ucsd.edu/ProteoSAFe/status.jsp?task=73da384ea02a4e8ca3edd82649e540c3. The MSMLS library can be accessed via the following link: https://gnps.ucsd.edu/ProteoSAFe/gnpslibrary.jsp?library=GNPS-MSMLS.

## 3. Results and Discussion

Proof-of-concept results were obtained using the MSMS-Chooser workflow on a 96-well plate containing 87 chemical standards. Positive and negative mode MS/MS spectra were collected using two separate injections into a flow of solvent. The method was 0.3 min in length providing enough time for the chemical standard to be detected and the line purged of any remaining material, as depicted in the chromatograms of two illustrative chemicals in positive and negative mode (**Figure 2A** and **Figure 2D**, respectively). The flow injection method was used for throughput data acquisition (~2 hrs for a 96-well plate), opposed to performing chromatographic separation; however, any method for MS/MS data acquisition is compatible with the MSMS-Chooser workflow.

**Figure 2.**
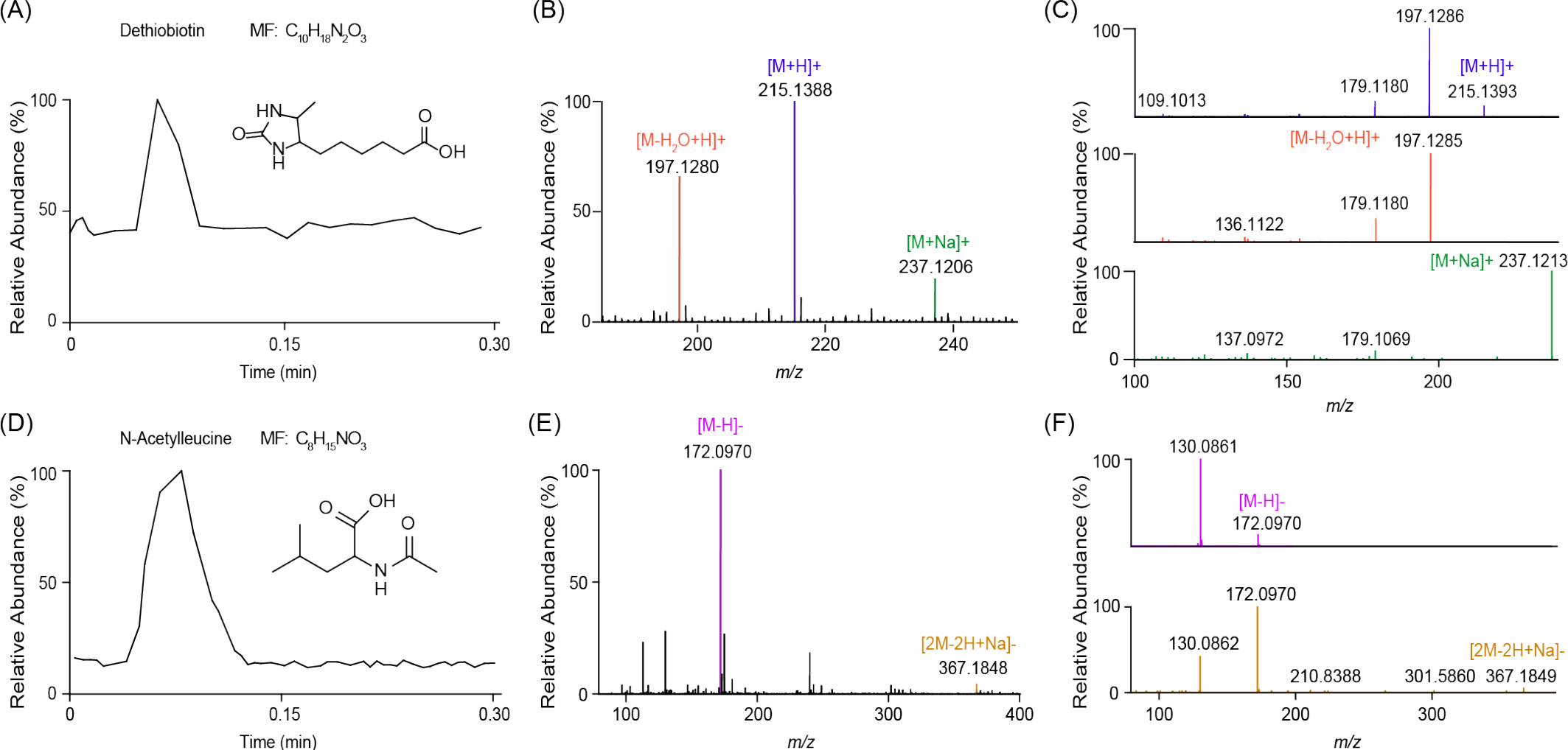
(A) Illustrative base peak chromatogram, (B) average MS spectrum, and (C) MS/MS spectra selected by MSMS-Chooser for dethiobiotin in the positive ionization mode. Peaks in the MS are colored corresponding to their MS/MS spectra. (D) Illustrative base peak chromatogram, (E) average MS spectrum, and (F) MS/MS spectra selected by MSMS-Chooser for N-acetylleucine in the negative ionization mode. Peaks in the MS are colored corresponding to their MS/MS spectra.

The data obtained were processed using the MSMS-Chooser workflow in GNPS to identify predesignated adducts calculated from the theoretical monoisotopic *m/z* within a mass error of 10 ppm. The resulting file from MSMS-Chooser indicates the scan number of the MS/MS spectrum and associated adduct for each chemical standard if detected. Multiple adducts were detected for some of the chemical standards which are formed during the electrospray ionization process under the experimental conditions (*e.g.* salinity, pH, and concentration). For example, the analysis of dethiobiotin in the positive ionization mode resulted in the detection of the protonated ([M+H]^+^), sodiated ([M+Na]^+^), and singly dehydrated species ([M−H_2_O+H]^+^), **Fig 2B**, and their corresponding MS/MS spectra selected by MSMS-Chooser (**Fig 2C**). Similarly, *N*-acetylleucine was detected as a deprotonated ([M−H]^−^) ion and as a [2M−2H+Na]^−^ ion in the negative ionization mode (**Fig 2D**), and MSMS-Chooser selected the corresponding MS/MS spectra (**Fig 2F**).

63 of the 87 chemical standards were detected upon manual inspection of the dataset. From the chemical standards detected, multiple hours of manual inspection found 73 MS/MS scans in positive mode and 53 MS/MS scans in negative mode (**Table 1**). MSMS-Chooser was able to correctly identify a total of 70 MS/MS scans in positive mode and 50 MS/MS scans in negative mode in minutes without user input. MSMS-Chooser correctly selected 99.2% of MS/MS spectra, excluding five MS/MS spectra not selected due to their mass error exceeding the 10 ppm tolerance. The adducts detected from this illustrative set of chemical standards were tabulated and plotted in **Figure 3A**. The predominant adducts in positive mode were [M+H]^+^ and [M+Na]^+^; and [M−H]^−^ in negative mode. The high levels of sodiated adducts observed, **Figure 3B**, is likely related to the number of sugars (*i.e.* Organooxygen compounds) included in the test dataset.

**Table 1.** Tabulated results comparing MSMS-Chooser and manual inspection of the data.

| Ionization Mode | Number of MS/MS spectra, Manual Inspection | Number of MS/MS spectra, MS-MS Chooser<sup>a</sup> | MS-MS Chooser Accuracy (%) | MS-MS Chooser Accuracy (%)<sup>b</sup> |
| --- | --- | --- | --- | --- |
| Positive | 73 | 70 | 95.9 | 98.6 |
| Negative | 53 | 50 | 94.3 | 100 |
| Total | 126 | 120 | 95.2 | 99.2 |

<sup>a</sup> MSMS-Chooser correctly identified 70 MS/MS spectra and incorrectly identified one MS/MS spectra in positive mode upon manual inspection which as removed for subsequent calculations.
<sup>b</sup> Manual inspection found 2 positive mode and 3 negative mode spectra that were not chosen by MSMS-Chooser. These spectra were excluded in the accuracy calculation as their mass error was greater than the 10 ppm mass error limit (an adjustable parameter) used in MSMS-Chooser.

**Figure 3.**
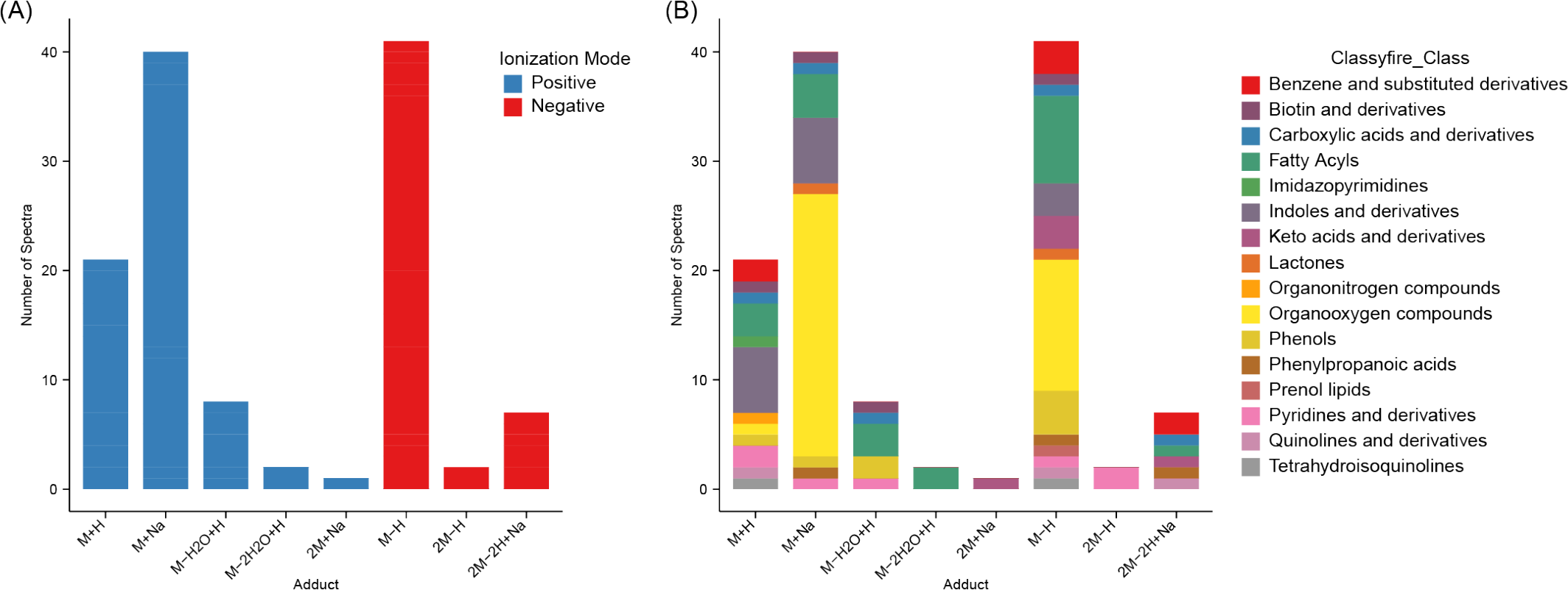
Adducts versus the number of MS/MS spectra selected by MSMS-Chooser in the proof-of-concept set of chemical standards, (A) colored by ionization mode and (B) colored by chemical classification, class level, obtained via Classyfire.

The MSMS-Chooser output provides users the ability to query their MS/MS spectra against all publicly available tandem MS datasets in GNPS via the tool MASST. The MASST results will indicate the datasets in which the query spectra had been previously detected. **Figure S1** depicts the MASST result of biotin, one of the chemical standards run in the MSMS-Chooser proof of concept dataset. Biotin was shown to be found in 16 GNPS datasets, comprised of microbial, humans, and food sample types.

## 4. Conclusion

The MSMS-Chooser workflow provides users with a web-based platform and instrument protocols for the rapid and systematic generation of MS/MS reference spectra. In our proof-of-concept results, MSMS-Chooser correctly identified 120 MS/MS spectra from 87 chemical standards in the positive and negative mode in minutes and output a file ready for the GNPS MS/MS spectral library. MSMS-Chooser was 99.2% accurate in selecting MS/MS spectra compared to manual evaluation (inspection of results is recommended). One strength of the workflow is the consideration of all possible adducts. Not all chemicals have the same predilection to ionization via one adduct (*e.g.* protonation or deprotonation). MSMS-Chooser assists in building MS/MS spectral libraries by considering multiple adducts that may be encountered using different electrospray ionization conditions. Any method of generating MS/MS (specifically product ion spectra) via different collisional activation methods is compatible with the GNPS spectral library. The current protocols do not capture collision energy but it is possible to do so; however, such information is not currently utilized in GNPS and therefore has limited utility. The variability in MS/MS spectra is best captured via the contribution of spectra of the same chemical collected on multiple instruments and multiple conditions. Contribution of all MS/MS spectra is highly encouraged.

MSMS-Chooser will aid the community of scientists who use GNPS (or the associated GNPS knowledge base) by increasing the number of MS/MS reference spectra in the public domain, improving annotation rates in untargeted metabolomics experiments. We demonstrated the ability to readily determine the location in which the chemical standards MS/MS spectrum selected via MSMS-Chooser were found using MASST.^11^ Any additional MS/MS spectra uploaded to the GNPS MS/MS spectral library will be included in the periodic living data analysis in GNPS. Data submitters will be emailed when new chemical annotations are found in their MassIVE deposited data. Further, additional MS/MS library spectra will extend the depth of annotation in ReDU,^12^ a recently developed web-enabled tool to reuse public MS/MS data. Lastly, the providence of the original data and MSMS-Chooser template is retained in MassIVE and available for re-analysis in a systematic and automated fashion.

## Acknowledgments

The authors would like to thank Julia M. Gauglitz, Ph.D. and Daniel Petras Ph.D. for insightful discussion. AKJ thanks the American Society for Mass Spectrometry for the Postdoctoral Career Development Award. The authors acknowledge funding by the Office of Naval Research Multidisciplinary University Research Initiative Award, (No. N00014-15-1-2809) and the Center for Microbiome Innovation. MC and PCD were supported by NSF grant IOS-1656475.

## Supplemental Information

**Figure S1.**
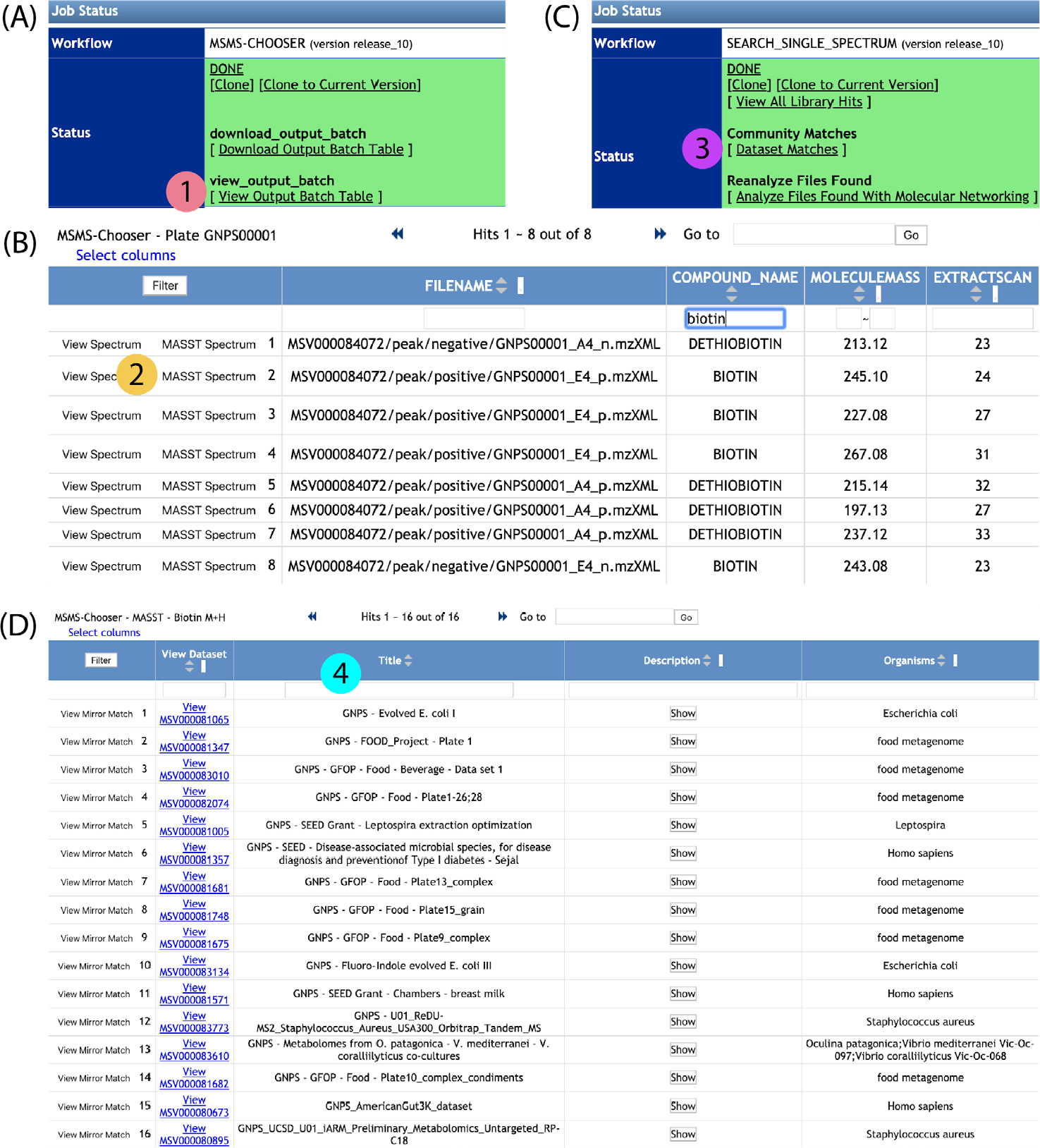
Overview of launching a MASST search from the MSMS-Chooser workflow results page. (A) MSMS-Chooser results page, (1) click on the “View Output Batch Table”. (B) View output batch page, (2) clicking on the “MASST Spectrum” button will launch a MASST search (*e.g.* biotin). (C) MASST search results page (https://gnps.ucsd.edu/ProteoSAFe/status.jsp?task=39cd886540c147e9a8a618b275e5f541), (3) click on the “Dataset Matches”. (D) Datasets in which biotin (an example) was detected using MASST, (4) the title and dataset accession link can be used to further investigate the data.

**Table S1.**
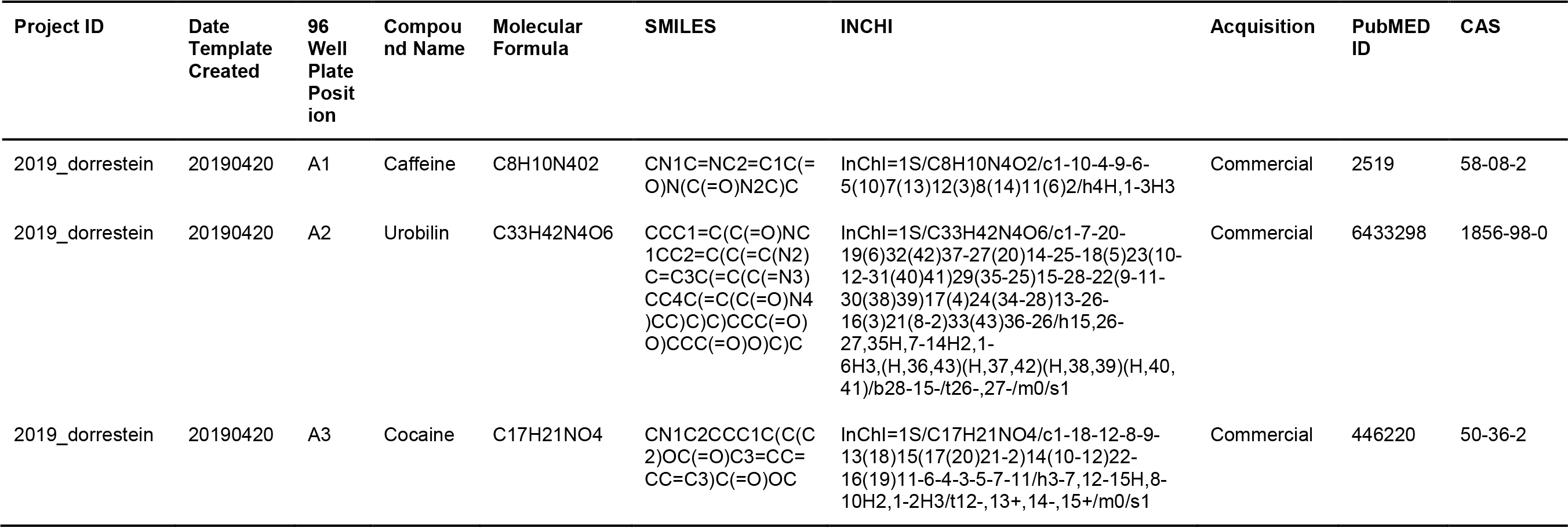
The minimum chemical information required for the MSMS-Chooser workflow which is found in the MSMS-Chooser template. Caffeine, urobilin, and cocaine are used as examples.

